# Tools enabling flexible approaches to high-resolution subtomogram averaging

**DOI:** 10.1101/2021.01.31.428990

**Authors:** Alister Burt, Lorenzo Gaifas, Tom Dendooven, Irina Gutsche

## Abstract

Cryo-electron tomography and subtomogram averaging are increasingly used for macromolecular structure determination *in situ*. Here we introduce a set of computational tools and resources designed to enable flexible approaches to subtomogram averaging. In particular, our tools simplify metadata handling, increase automation, and interface the *Dynamo* software package with the *Warp-Relion-M* pipeline. We provide a framework for *ab initio* and geometrical approaches to subtomogram averaging combining tools from these packages. We illustrate the power of working within the framework enabled by our developments by applying it to *EMPIAR-10164*, a publicly available dataset containing immature HIV-1 virus-like particles, and a challenging *in situ* dataset containing chemosensory arrays in bacterial minicells. Additionally, we establish an open and collaborative online platform for sharing knowledge and tools related to cryo-electron tomography data processing. To this platform, we contribute a comprehensive guide to obtaining state-of-the-art results from *EMPIAR-10164*.

## Introduction

Cryo-electron tomography (cryo-ET) is an imaging technique rapidly gaining popularity for the direct visualisation of unique biological objects in 3D. Repeating structural motifs present in cryo-ET data can be reconstructed at higher resolution by subtomogram averaging (STA), allowing for the possibility of studying macromolecular structure *in situ* (Schur, 2019). STA has developed alongside single particle analysis (SPA) in cryo-electron microscopy (cryo-EM), a technique which has benefited significantly in the last ten years from advances in both electron detection hardware and image processing software (Bai et al., 2015; Nakane et al., 2020).

State-of-the-art subtomogram averaging workflows often co-opt tools and adapt ideas from single particle analysis for tomography and subtomogram averaging (Sanchez et al., 2020; Turoňová et al., 2020). An unfortunate consequence of this side-by-side development is a somewhat fragmented software ecosystem which employs various file formats and metadata conventions (Leigh et al., 2019). Despite the advent of complete or near complete integrated solutions for cryo-ET and STA (Chen et al., 2019; Himes and Zhang, 2018; Tegunov et al., 2021), employing optimal methodology for a given dataset often requires the creative combination of different approaches which may not all be present within a single integrated pipeline. For those new to cryo-ET, the burden of interfacing many different software packages in a complex workflow often represents a barrier to the testing of alternative approaches.

Image processing for STA from cryo-ET data is a complex process which can broadly be divided into three blocks (Figure 1). The first, ‘preprocessing’, transforms experimental 2D micrographs into 3D reconstructions of an imaged region. The second, ‘particle picking’, generates putative positions and orientations for objects of interest within each volume as well as initial reference(s) for subsequent refinement. Finally, the ‘refinement’ block is concerned with the optimisation of reconstruction(s) from imaging data associated with each particle position.

**Figure 1:**
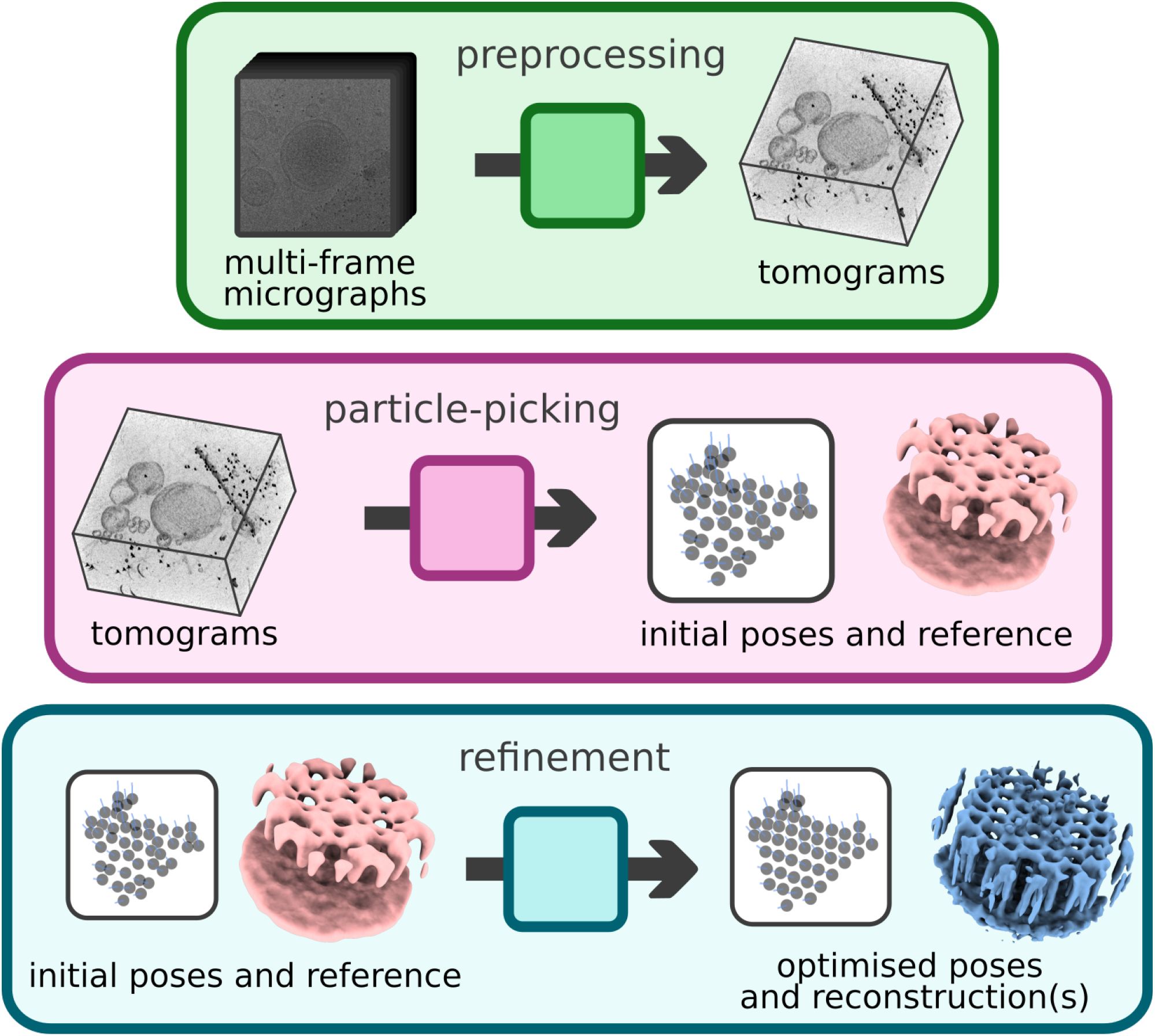
Cryo-ET data processing can be roughly divided into three blocks. Preprocessing (top) transforms raw experimental data, usually in the form of multi-frame micrographs, into 3D images called tomograms. Particle picking (middle) derives particle position and orientation information from tomograms as well as an initial reference for subsequent refinement. Refinement (bottom) optimises (multiple) 3D reconstructions from input data.

The preprocessing block encompasses all steps in the generation of tomograms from experimental data. Experimental cryo-ET data is usually acquired as a set of multi-frame 2D micrographs, one per tilt angle in a tilt-series. Typically, this block includes per-tilt inter-frame motion estimation and correction, contrast transfer function (CTF) estimation, tilt-series alignment and 3D reconstruction (Tegunov and Cramer, 2019). Tilt-series alignment is the least automated of these steps, with significant time often spent optimising alignments in an attempt to produce more accurate reconstructions.

Particle picking can be somewhat entangled with the process of initial model generation, with some particle picking methods requiring an initial model and others being reference-free. Exhaustive, reference-based template matching approaches are widely accepted as an imperfect solution for particle picking due to the computational overhead and the need for subsequent dataset cleaning (Basanta et al., 2020; Schaffer et al., 2019). Care must also be taken to avoid model bias. Particle picking methods based on deep-learning have been proposed, which may help to address some of these limitations (Chen et al., 2017; Moebel et al., 2021). Another class of particle picking methods derive putative particle positions and orientations from a 3D model of a supporting geometry, usually generated from minimal manual annotations (Castaño-Díez et al., 2017). These methods can be used to impose prior knowledge about how particle poses relate to the supporting geometry during refinement. Employing such *a priori* knowledge is advantageous, reducing both the computational burden of global searches and the likelihood of ending up in incorrect local minima during refinement. These advantages come at the expense of extra time spent on manual annotation and management of more metadata.

Refinement has typically meant the optimisation of particle poses within fixed 3D reconstructions (Wan and Briggs, 2016). More recently, this block has been extended to include procedures for the optimisation of many geometrical and electron-optical parameters, improving the resolutions attainable by STA (Bartesaghi et al., 2012; Chen et al., 2019; Himes and Zhang, 2018; Tegunov et al., 2021).

Since the advent of methods for 3D contrast transfer function (CTF) correction (Turoňová et al., 2017) and *a posteriori* improvement of an existing set of alignment parameters (Bartesaghi et al., 2012; Chen et al., 2019; Himes and Zhang, 2018; Tegunov et al., 2021), the potential for studying macromolecular structure at intermediate resolution (3-5 Å) by STA is becoming a reality. Despite these advances, there remain relatively few examples of STA reconstructions in this resolution regime in the Electron Microscopy Data Bank (EMDB), indicating that this work remains non-trivial (Figure 1-figure supplement 1).

*Dynamo* is a flexible, extensible software environment for subtomogram averaging, providing a variety of powerful tools for data management, annotation and analysis (Castaño-Díez et al., 2012). The *Warp-RELION-M* pipeline is an integrated solution for cryo-ET and SPA, yielding the highest resolution reconstructions from cryo-ET data thus far (Tegunov et al., 2021). In the materials and methods section, we describe tools designed to facilitate certain computational aspects of cryo-ET by simplifying metadata handling, increasing automation and interfacing *Dynamo* with the *Warp-RELION-M* pipeline.

In the results and discussion section, we show that combination of these tools enables working within a flexible framework for *ab initio* and geometrical approaches to subtomogram averaging. We illustrate the power of working within such a framework on two datasets. We also provide a collaborative, online platform for the community to share knowledge about cryo-ET data processing, to which we contribute a complete guide for achieving state-of-the-art results on a five tomogram subset of *EMPIAR-10164*.

## Materials and Methods

### Integrating on-the-fly tilt-series alignment and data management into the Warp preprocessing pipeline

Tomogram reconstruction within the *Warp* preprocessing pipeline currently requires the use of an external software package, *IMOD* (Kremer et al., 1996), for tilt-series alignment. In the absence of a fully integrated solution in the currently available version of *Warp* (1.0.9), we developed *autoalign_dynamo*, a package for automating fiducial-based tilt-series alignment using *Dynamo* and *IMOD. dautoalign4warp* provides a simple command line interface for on-the-fly alignment of tiltseries. The program proceeds by dynamic generation of a *Dynamo* tilt-series alignment workflow with appropriate parameters derived from minimal user input. The workflow is executed, generating a set of refined fiducial positions. Refined fiducial positions are converted using *dms2mod* (provided by *autoalign_dynamo*) and used to generate tilt-series alignment parameters using the *IMOD* program *tiltalign*. Key *tiltalign* parameters fixed within this procedure are: solve for one tilt axis for the entire tilt-series, fix tilt-angles at their nominal values, use robust fitting for parameter estimation. All parameters can be seen and modified in the *tiltalign_wrapper* script. All data is subsequently organised such that alignments can be imported directly into Warp for tomogram reconstruction, requiring no further user input. The base function *autoalign* which wraps *Dynamo* tilt-series alignment functionality is provided for those who wish to apply this procedure outside of the *Warp* pipeline. A function *warp2catalogue* serves to set up a database called a *Dynamo* catalogue for tomograms reconstructed in *Warp*. The catalogue is set up such that all visualisation operations make use of a deconvolved reconstruction, filtered for optimal visualisation, whilst all particle extraction operations make use of the corresponding unfiltered volume.

Dynamo tilt-series alignment workflows were released recently and as such are not yet described in the literature, although details and the source code are available in the public domain (Castaño-Díez, 2019). To facilitate the readers understanding, we provide a brief overview of the alignment algorithm, noting that it was not implemented by us. A binary, synthetic template of a fiducial marker is generated based on user input and used to detect candidate fiducial positions within a tilt-series by cross-correlation. 300 sub-images are extracted from the tilt-series at the peaks of the resulting cross-correlation matrix and averaged to produce a template for detecting fiducial markers in the data. The template is used to detect initial fiducial positions in the tilt-series by cross-correlation. Cross-correlation peaks are analysed and those observations not meeting a minimum degree of rotational symmetry are discarded. Observations are indexed to link observations of the same fiducial marker in multiple micrographs by pairwise cross-correlation of the observations between neighbouring micrographs. The longest ‘trails’ of linked observations are used to generate an initial 3D model of fiducial positions. The 3D model is iteratively reindexed by reprojection of fiducial positions against the tilt-series, adding observations which fall within a distance threshold to the set of observations to be included for further refinement. Tilt-images lacking at this point are reintegrated by a procedure comparing the reprojection of the current model with the cloud of initial observations found on that micrograph. The number of observations is maximised by the reintegration of missing fiducial markers from other images using the same procedure. The projection model is iteratively refined before the positions of fiducial markers are independently iteratively refined against an average of all observations of that marker. The final set of refined markers is pruned according to the root mean square deviation (RMSD) between measured fiducial position and the reprojected position of the 3D model.

### Python package for Euler angle conversions

Euler angles are used by many cryo-EM software packages to describe the orientation of a rigid body with respect to a fixed coordinate system. *eulerangles* is a self-contained Python package which provides a simple application programming interface (API) for the batch conversion of Euler angles into rotation matrices, rotation matrices into Euler angles and interconversion of Euler angles defined according to different conventions. Conversions can take place between all possible definitions of Euler angles in a right-handed coordinate system and the implementation is vectorised, allowing it to handle large datasets efficiently. Conventions for Euler angles from *Dynamo* and *Warp-RELION-M* are provided in the library itself, along with a simple mechanism for the definition of conventions from other software packages.

### Python packages for metadata handling

We have developed two self-contained Python packages *starfile* and *dynamotable* to facilitate handling the primary metadata systems of *RELION* (Scheres, 2012) *a*nd *Dynamo* (Castaño-Díez et al., 2012). These packages provide input/output functionality via a simple API. Data is exposed as *pandas* DataFrame objects, effectively interfacing these metadata systems directly with the scientific Python (Virtanen et al., 2020) ecosystem.

### Integration of data collected in Tomography 5 as multi-frame micrographs into the Warp preprocessing pipeline

In the *Warp* cryo-ET preprocessing pipeline, only metadata from the *SerialEM* data collection program in the form of mdoc files is currently supported. Thermo Scientific *Tomography 5* (Tomo 5) is the official solution provided with Thermo Scientific microscopes for electron tomography experiments (ThermoFisher Scientific, 2020). We provide a small command line tool *mdocspoofer* for the generation of these metadata files for a directory containing multi-frame micrograph files of the form *<basename>_<count>[<tilt_angle>]_fractions.mrc*, as generated by *Tomography 5*. This tool enables use of the *Warp* preprocessing pipeline for data collected in *Tomography 5*.

### Interfacing Dynamo and the Warp-RELION-M pipeline

The recommended procedure for particle picking in the *Warp-RELION-M* pipeline is exhaustive, reference-based template matching. For initial particle pose optimisation, *RELION* subtomogram averaging is integrated into the pipeline. Alternative approaches to particle picking, pose optimisation and classification may provide advantages over this workflow. We provide *dynamo2m*, a set of tools which create a bidirectional interface between *Dynamo* and the *Warp-RELION-M* pipeline. *dynamo2warp* allows for particle position and orientation data in Dynamo to be used for particle extraction in *Warp* and subsequent alignment in *RELION* and *M*. *warp2dynamo* provides a route to using particles extracted in *Warp* within *Dynamo*. Because *RELION* modified its metadata system with the release of *RELION 3.1, relion_star_downgrade* is provided to enable extracting particles refined using *RELION* version 3.1 or higher in *Warp* 1.0.9.

## Results and Discussion

### Metadata handling

The ability to test alternative approaches and rapidly iterate is key to optimising complex data analysis workflows like cryo-ET and STA. Converting between different Euler angle conventions is often a pain point for those wishing to interface different software packages due to the abundance of ambiguities in their interpretation (Urzhumtseva and Urzhumtsev, 2019). The *eulerangles* package simplifies both interfacing pieces of software which interpret Euler angles according to differing conventions and making use of Euler angles for custom analyses or visualisations.

Implementing custom input/output functionality when attempting to work with a variety of non-standard metadata presents a barrier to entry for those wishing to interactively explore their data or implement custom analysis routines. The Python packages *starfile* and *dynamotable* remove this barrier for the working with *RELION* and *Dynamo*, providing a simple interface between their respective metadata systems and a wealth of data analysis infrastructure, visualisation tools and educational resources (Harris et al., 2020; Paszke et al., 2019).

Tight integration with the scientific Python ecosystem encourages exploratory data analysis and simplifies the development of any custom tooling, as is often necessary in cryo-ET. As an example of the benefits derived from working within this ecosystem, we combine our *starfile* and *eulerangles* packages with the packages *mrcfile* (Burnley et al., 2017) (for reading and writing MRC format image files) and *napari* (Sofroniew et al., 2021) (a fast, powerful multidimensional data visualisation library) to create a visualisation of particle positions, their orientations and tomograms in 3D (Figure 2). By combining these packages, creating such a visualisation from scratch can be achieved in less than 25 lines of code (Figure 2-figure supplement 1).

**Figure 2:**
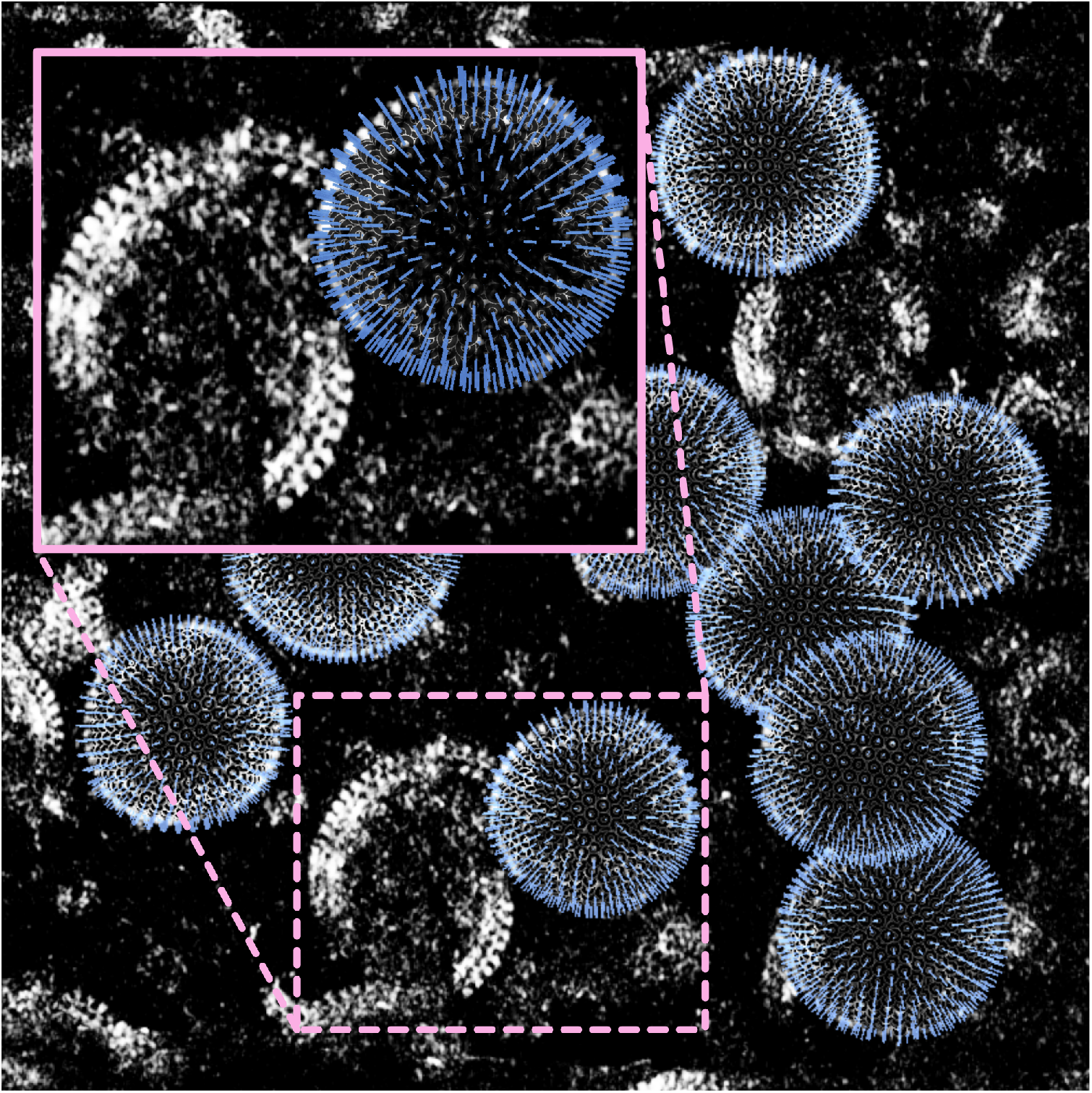
Modular tools designed to integrate with the scientific Python ecosystem simplify the creation of custom visualisations in *napari*. Particle positions and orientations, easily accessed and manipulated using *starfile* and *eulerangles*, are rendered in 3D with the imaging data from which they were derived.

### The *Dynamo - Warp-RELION-M* interface

In this section, we discuss and illustrate the benefits provided by the bidirectional *Dynamo - Warp-RELION-M* interface.

#### Automation of tilt-series alignment in *Warp-RELION-M*

Optimisation of data collection strategies has significantly increased the throughput of cryo-ET data collection in recent years, with datasets now routinely exceeding 100 tomograms. As dataset sizes increase, so does the need for accurate, automated solutions to each and every step of complex data analysis workflows. Accurate tilt-series alignment is often achieved using semiautomated procedures which quickly become tedious when faced with a large dataset. Providing the automated, robust fiducial-based tilt-series alignment procedures from *Dynamo* as an on-the-fly solution integrated into the *Warp* preprocessing pipeline using *autoalign_dynamo* greatly simplifies the generation of accurate 3D reconstructions for downstream data analysis. We provide no integrated solution for aligning tilt-series lacking exogenous fiducial markers, such as those from samples prepared by focused-ion beam (FIB) milling, although procedures exist in other software packages (Chen et al., 2019). As new procedures are developed, their integration into on-the-fly data processing workflows will be important for optimal use of both microscope and researcher time.

#### Enabling geometrical approaches to subtomogram averaging in *Warp-RELION-M*

*warp2catalogue* and *dynamo2warp* enable the use of geometrical approaches to subtomogram averaging to users of the *Warp-RELION-M* pipeline (Figure 3). The catalogue system in *Dynamo* and the interactive 3D modelling tools in the *dtmslice* viewer make it easy to generate models of supporting geometries and derive particle positions and orientations for an entire dataset from minimal manual annotations (Castaño-Díez et al., 2017). Examples of easy-to-annotate supporting geometries from which positions can be easily derived are vesicles (Figure 3a), arbitrarily shaped membrane surfaces (Figure 3b), filaments and crystals. *warp2catalogue* simplifies the setup of a Dynamo catalogue for *Warp-RELION-M* users: the user annotates volumes filtered to aid visualisation, whereas particle extraction operations performed from the catalogue use the corresponding unfiltered data. This automation simplifies the use of an optimal workflow without unnecessary cognitive burden for the user.

**Figure 3:**
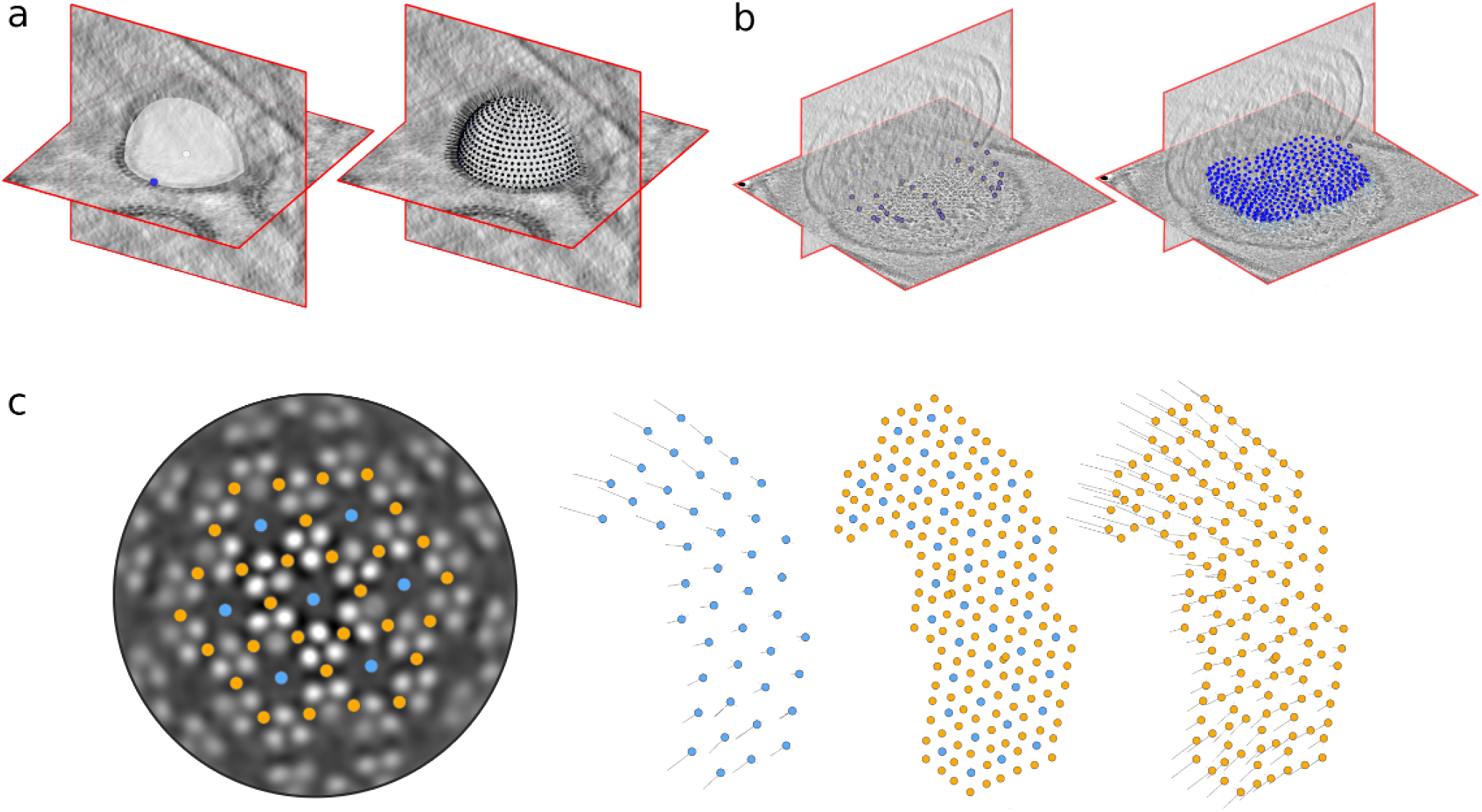
Geometrical approaches to subtomogram averaging. Particle positions and orientations are generated from minimal manual annotations for (a) a spherical vesicle and (b) an arbitrarily shaped surface. (c) different positions in a chemosensory array (EMD-10160) are annotated in blue and orange. A set of particle poses from a subtomogram averaging experiment (blue) are used to derive the positions and orientations of the orange particles in a ‘subboxing’ procedure.

*Dynamo* also provides *dpktbl.subbox.tableOnTable* for ‘subboxing’, deriving particle positions and orientations which are geometrically related to an existing set of positions and orientations. (Figure 3c). This procedure is often useful in cryo-ET for focussing analysis on subunits of a large complex after an initial consensus refinement (Ertel et al., 2017). *dynamo2warp* provides a simple mechanism for the integration of these powerful tools into the *Warp-RELION-M* pipeline.

#### Providing access to alternative refinement and classification procedures

*RELION* implements a Bayesian approach to the refinement of one or many 3D volumes from cryo-EM data, called maximum *a posteriori* refinement (Scheres, 2012). Originally designed for single particle cryo-EM data and more recently extended to work with cryo-ET data (Bharat et al., 2015), *RELION* provides solutions for the ‘refinement’ block of the cryo-ET pipeline. The package features an easy-to-use graphical user interface (GUI) and is already in use in a large number of cryo-EM labs. Rather than a data processing pipeline, *Dynamo* is a collection of powerful tools for working with subtomogram averaging data. With an emphasis on customisation, *Dynamo* often exposes the user to large numbers of parameters in fully featured GUIs.

By interfacing *Dynamo* with the *Warp-RELION-M* pipeline in the ‘refinement’ block, we provide *Warp-RELION-M* users access to highly customisable subtomogram averaging and multireference classification procedures, Principal Component Analysis (PCA)-based classification and myriad data visualisation tools and analysis tools. *Dynamo* users gain a route to making use of the CTF estimation and particle reconstruction tools in *Warp*, subtomogram averaging and classification in *RELION* and the multi-particle refinement procedure implemented in *M*.

As an example of a possible benefit of increased flexibility in subtomogram averaging workflows, we compare the options for masking between *Dynamo* and *RELION*. In *Dynamo*, a user may choose to provide separate masks for alignment, classification and Fourier shell correlation (FSC) to be used during an iterative alignment project. In *RELION*, the same mask is used for all three. Creative combination of different masks can allow for excluding a membrane from alignment for a membrane protein which would otherwise require using masks with extremely soft edges to avoid FSC artefacts. Such a soft mask would necessarily include the membrane or exclude membrane-proximal protein density.

### A framework for flexible ab initio and geometrical approaches to subtomogram averaging

Each subtomogram averaging dataset is different and analysis presents its own unique challenges. The provided tools were designed to simplify cryo-ET data processing where possible and empower researchers with access to a vast array of alternative tooling where it may provide benefit. Combined, these tools enable working within a powerful, flexible framework for *ab initio* and geometrical approaches to subtomogram averaging, combining *Warp-RELION-M* with *Dynamo* (Figure 4).

**Figure 4:**
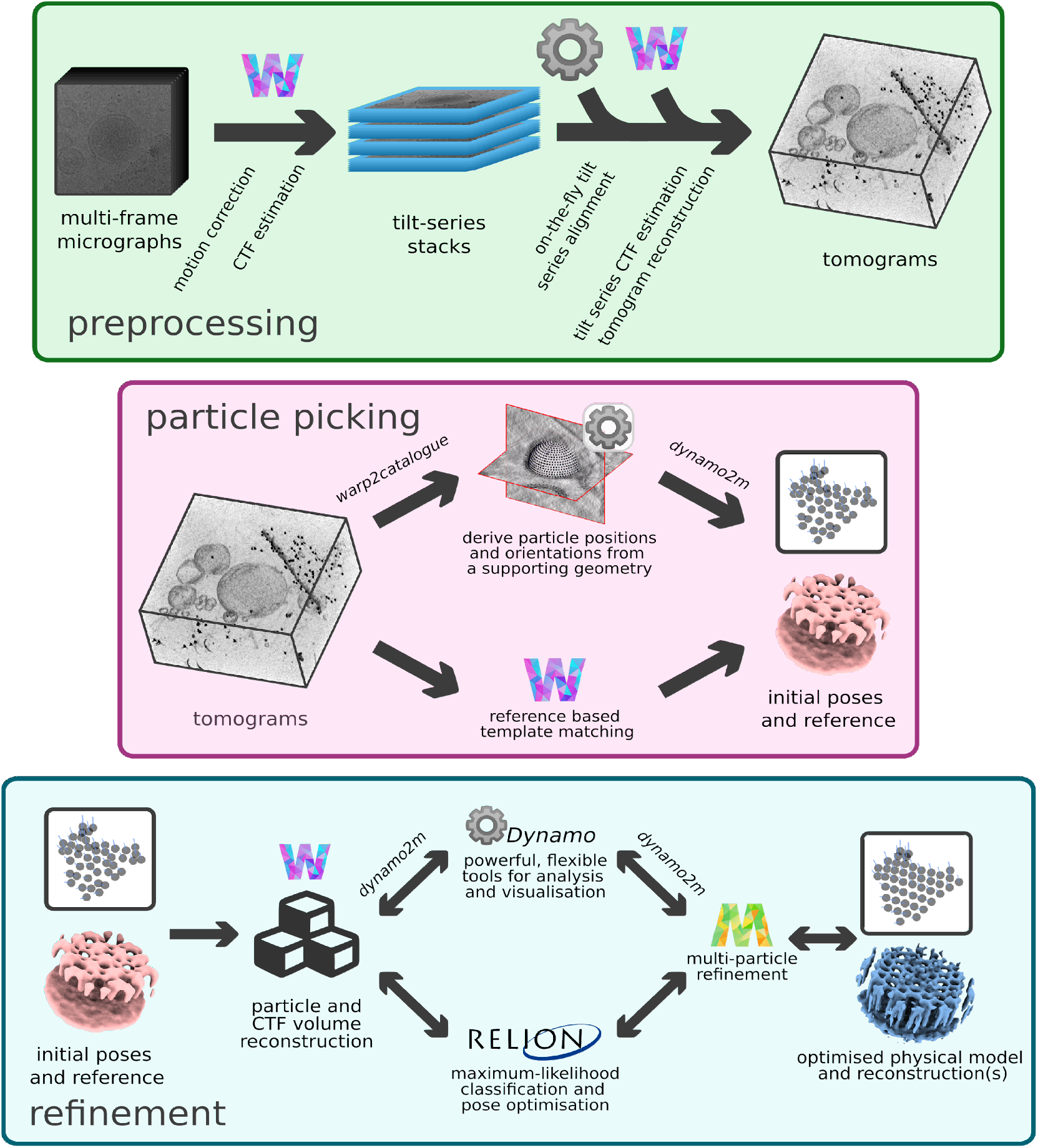
*autoalign_dynamo* and *dynamo2m* enable working within a powerful, flexible framework for *ab initio* and geometrical approaches to subtomogram averaging. The cryo-ET preprocessing pipeline in *Warp* (top) is extended to include on-the-fly tilt-series alignment in *Dynamo*. Tools for deriving particle positions and orientations from a 3D model of a supporting geometry are integrated into the particle picking block. In the refinement block, users of *Warp-RELION-M* and *Dynamo* can freely move between each package, choosing the tools most suited to the problems posed by their data.

#### Application to the HIV-1 CA-SP1 hexamer (EMPIAR-10164)

We illustrate the power of working within this framework on an example dataset by reprocessing a five tilt-series subset of *EMPIAR-10164* containing immature HIV-1 dMACANC viruslike particles (VLPs). This dataset was contributed to the community by the Briggs group and this subset has previously been used to benchmark *NovaCTF* (Turoňová et al., 2017) and *Warp* (Tegunov and Cramer, 2019), resulting in 3.9 Å and 3.8 Å reconstructions respectively.

Initial motion correction and initial CTF estimation were performed in *Warp* with respective spatiotemporal model resolutions of (1, 1, 8) and (2, 2, 1). Tilt-series were automatically aligned using *autoalign_dynamo* before CTF estimation and tomogram reconstruction at 10 Å/px in *Warp*. VLPs were annotated in *Dynamo* and initial estimates of positions and orientations were generated with an inter-particle distance of 45 Å, oversampled relative to the ~75 Å lattice spacing seen in the tomograms. 500 particles were extracted and averaged in *Dynamo* to produce a coarse template. The same data was subject to subtomogram averaging against this template in *Dynamo* without imposing symmetry during refinement. The resulting average contained a hexagonal lattice. To obtain an initial model, the 6-fold axis of the lattice was aligned to the z-axis and the resulting volume was symmetrised 6-fold in *Dynamo*. 28516 particles were extracted from annotated VLPs and aligned against the initial model for one iteration with a limited angular search range in *Dynamo*. The resulting particle positions and orientations, visualised in *Dynamo*, formed regular lattices on the surfaces of each VLP with some less regular areas. Using simple MATLAB scripts, only the 19810 particles with three or more neighbours at the expected inter-particle distance were retained for subsequent analysis. Using *dynamo2m*, metadata were converted to enable working in *Warp-RELION-M*. Particle extractions were carried out in Warp and 3D refinements in *RELION 3.1*, starting with local angular searches of ±15°. Resolution estimates were measured by FSC between independent reconstructions from random half-sets of the data using *relion_postprocess*. Particles were (i) extracted at 5 Å/px and refined (unmasked) to 10 Å resolution, (ii) extracted at 2.5 Å/px and refined (unmasked) to 5 Å resolution, (iii) extracted at 1.7 Å/px in *Warp* and refined (masked to include only the central hexamer) to 3.8 Å resolution. This intermediate result is indicative of the accuracy of fiducial-based alignment workflows in *Dynamo* and alignments in *RELION*. Seven rounds of multiparticle refinement were performed in *M* at 1.6 Å/px in a masked region containing only the central hexamer. The first 4 rounds of multi-particle refinement, optimising only for deformation parameters, yielded a 3.6 Å reconstruction. Three further rounds including optimisation of electron-optical aberration related parameters and tilt-frame alignments refined to a resolution of 3.4 Å (Figure 5).

**Figure 5:**
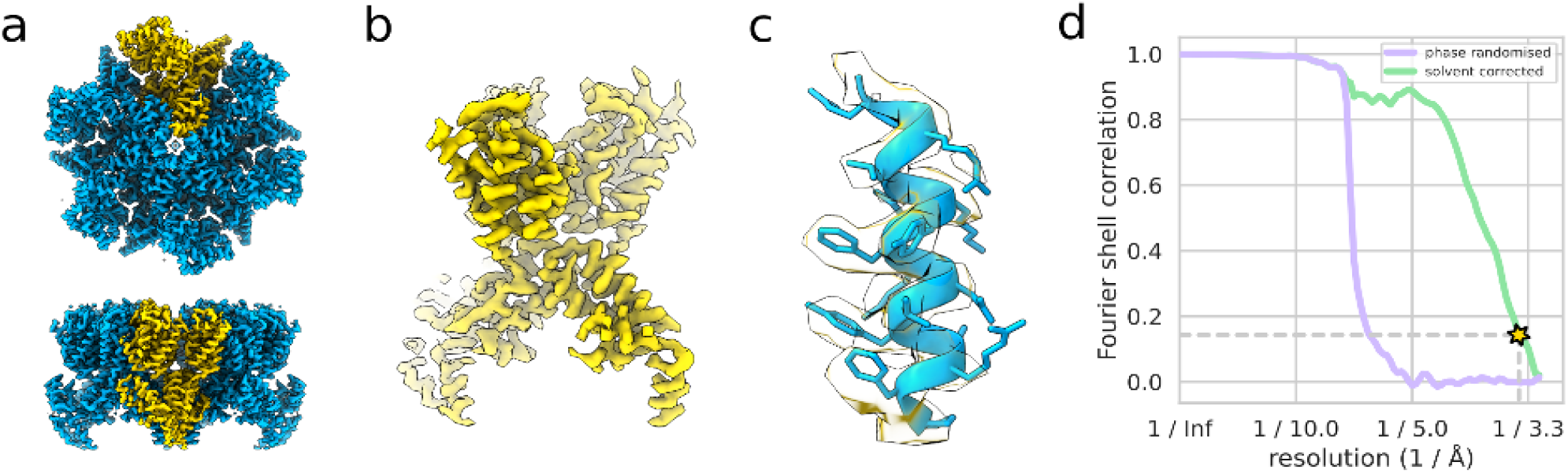
A (nominally) 3.4 Å reconstruction of the HIV-1 CA-SP1 hexamer from 19810 particles in 5 tiltseries of *EMPIAR-10164*. The whole hexamer with one monomer highlighted in yellow (a), one monomer (b) and the region of the map around helix 292-307 PDB 5L93 (fitted) (c) are shown. The gold-standard FSC curve (d) used to estimate the nominal resolution value is shown as well as the phase-randomised masked FSC used to validate the mask used for FSC calculation.

It should be noted that working within this framework readily yielded a 3.4 Å reconstruction from 19810 particles in five tomograms, without use of an external reference and assuming no prior knowledge about the complex. The combination of *Dynamo* tools and the *Warp-RELION-M* pipeline, enabled by the tools presented here, rendered this *ab initio*, geometrical approach a smooth, efficient process. An *ab initio* approach is not strictly necessary for this dataset, as demonstrated by *emClarity* (Himes and Zhang, 2018) and *Warp-RELION-M* (Tegunov et al., 2021). We however note that obtaining an initial model is often a challenging step in the first stages of a project. Thus, access to *ab initio* and geometrical approaches is provided as an alternative should the integrated template matching procedure prove inadequate. The final resolution of the reconstruction serves only to demonstrate the ability of *M* to further optimise image alignment and electron-optical parameters.

#### Application to the E. *coli* chemosensory array *in situ*

We are currently using an optimised minicell system (Burt et al., 2020) to investigate structural bases of the chemotactic response in motile bacteria. To illustrate the advantages of working within this framework, we choose to present intermediate results from a set of 109 minicell tomograms (Figure 6).

**Figure 6:**
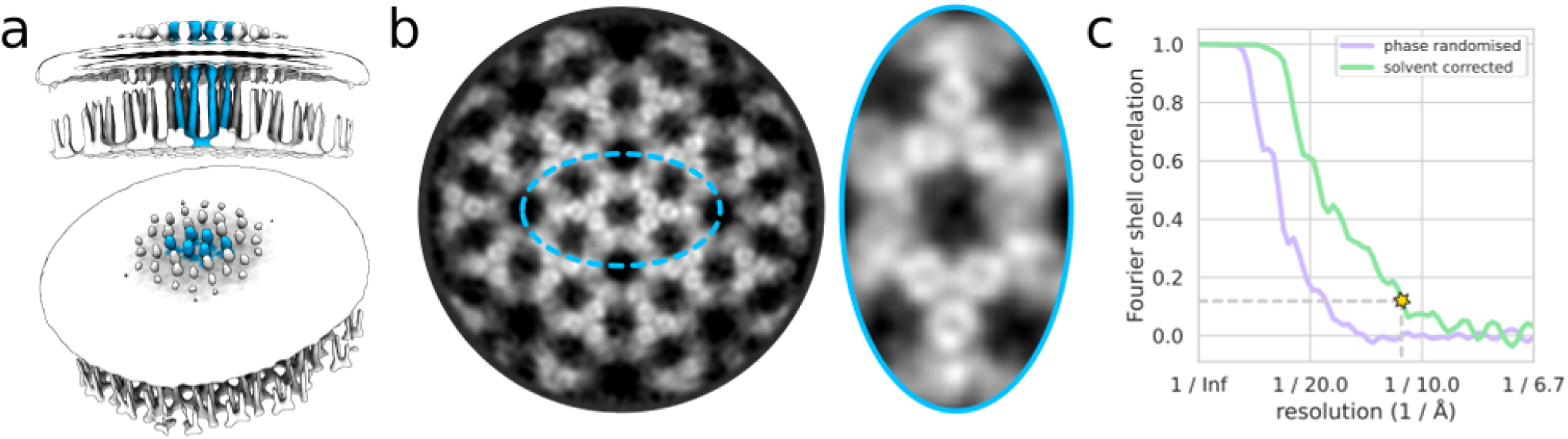
A (nominally) 11 Å reconstruction of the *in situ* chemosensory array from 17484 particles in 48 tomograms of *E. coli* minicells. The full reconstruction with a pixel size of 7.5 Å/px is shown in (a) with the membrane density superposed at a lower isosurface threshold. The quality of the density is visible in a slice through the map reconstructed with a pixel size of 2.24 Å/px and filtered to 11 Å (b - left). Holes are clearly visible in the densities of receptor dimers at this resolution (b - right). The gold-standard FSC curve (c) used to estimate the nominal resolution value is shown alongside the phase-randomised masked FSC curve used to validate the mask used for FSC calculation.

The chemotaxis-mediating molecular machinery forms large supramolecular arrays, a highly cooperative network of core signalling units (CSUs), in the bacterial inner membrane at the cell poles. The elongated nature of the CSU, the crowded cellular milieu and increased sample thickness complicate template matching approaches with this system, increasing the false-positive rate and impeding appropriate peak extraction. Instead, we opt for a seed-oversampling approach in which initial positions and orientations are distributed on *Dynamo* surface models, oversampling with respect to the expected inter-particle distance of ~120 Å. Particle positions were generated from the surface model with an inter-particle distance of 30 Å using *Dynamo*. Particles were extracted and subject to one round of subtomogram averaging in *Dynamo* with local angular searches of ±30°. Constraining the angles of particles relative to initial estimates from a supporting geometry ensures that particles within each array have orientational parameters coherent with our understanding of the system. Particles were extracted at 7.5 Å/px in *Warp* and imported into *Dynamo* using *warp2dynamo*.

A consensus refinement of these particles, centered on the three-fold symmetry axis of a p6-symmetric assembly, was performed in *Dynamo*, allowing local deviations in out-of-plane orientation, complete search of in-plane orientation and enforcing C3 symmetry during refinement. In the resulting average, the expected 3-fold symmetric arrangement of P4 domains in the structure was not readily observed. The resulting reconstruction had a nominal resolution of 18 Å, with the FSC curve indicating the presence of significant heterogeneity. The *Dynamo* tool *dpktbl.subbox.tableOnTable* was used to derive the positions and orientations of particles centered on six inter-trimer-of-dimer axes of the array. Duplicate particles were removed using *dpktbl.exclusionPerVolume* prior to a consensus refinement of these particles enforcing C2 symmetry. The resulting reconstruction had a nominal resolution of 19 Å. Attempts to disentangle structural heterogeneity using multireference approaches to 3D classification in both *Dynamo* and *RELION* failed to yield sensible results at this stage.

Multi-particle refinement in *M* yielded improvements when performed in a masked region encompassing a large array region of 400 Å diameter, resulting in a reconstruction with a nominal resolution of 11 Å. No improvement was seen when refinement was performed on smaller regions encompassing one to four CSUs, presumably due to insufficient signal. Work is underway to improve upon this intermediate result by classification and further refinement. However, these initial results are already quite promising, showing features such as a hole at the center of the four-helical bundle of a receptor dimer, not yet seen *in situ*.

### A platform for collaborative knowledge sharing in the cryo-ET community

Information about best practices and approaches to problems presented by cryo-ET data analysis is currently divided between the literature, mailing lists and the documentation of various pieces of software. We establish teamtomo.org as an open, collaborative platform for knowledge sharing within the growing cryo-ET community. The platform is designed from the ground-up as a collaborative endeavour with built-in mechanisms for easily integrating contributions.

Intended for sharing information about general data processing principles, problem solving approaches, short and long-form tutorials as well as providing links to relevant external material, the platform provides an overview of the field for interested parties, as well as specific practical and theoretical knowledge for those actively processing data. As an initial contribution, we provide a complete guide for the procedure we used to achieve the 3.4 Å reconstruction of the HIV-1 CA-SP1 hexamer, together with practical and theoretical tips of relevance for this and other datasets. Those wishing to share their expertise are encouraged to contribute.

## Conclusions

The tools and resources presented here increase automation within the *Warp* preprocessing pipeline, simplify metadata handling and interface two complementary cryo-ET data processing solutions. The provided interface enables working within a powerful, flexible framework for geometrical and *ab initio* approaches to subtomogram averaging which can yield high resolution results. Our tools increase the functionality available to users of both *Dynamo* and *Warp-RELION-M*, lowering the barrier to employing more customisable workflows for cryo-ET data processing which are often key to success. Finally, we hope that the existence of a collaborative, community-driven resource for sharing cryo-ET knowledge will encourage open science, improve the ability of the growing cryo-ET community to integrate newcomers and enable their use of cryo-ET to solve lifescience questions.

## Data availability

Data on immature HIV-1 dMACANC virus-like particles used for benchmarking of the pipeline was taken from EMPIAR-10164. Data on *E. coli* minicells will be available upon publication of the final reconstruction of the chemosensory array.

## Code availability

Source code for all tools and resources described here is available at:

- *autoalign_dynamo* - https://github.com/alisterburt/autoalign_dynamo
- *mdocspoofer* - https://github.com/alisterburt/mdocspoofer
- *starfile* - https://github.com/alisterburt/starfile
- *dynamotable* - https://github.com/alisterburt/dynamotable
- *eulerangles* - https://github.com/alisterburt/eulerangles

All Python packages are made available on the Python package index (PyPI).

## Acknowledgements

This work has received funding from a European Union’s Horizon 2020 research and innovation programme under grant agreement No 647784 to IG. We acknowledge Diamond Light Source for access and support of the cryo-EM facilities at the UK’s national Electron Bio-imaging Centre (eBIC), funded by the Wellcome Trust, MRC and BBRSC. Cryo-ET data acquisition has been supported by iNEXT, grant number 653706 (PID:2626 to IG), funded by the EU Horizon 2020 programme. AB is supported by a University Grenoble Alpes PhD fellowship and by a Fondation pour la Recherche Medicale (FRM) fellowship. LG is supported by a PhD fellowship from Grenoble Alliance for Integrated Structural and Cell Biology (GRAL, ANR-10-LABX-49-01) funded within the CBH Graduate School of the University Grenoble Alpes (ANR-17-EURE-0003).

## References

Bai X, McMullan G, Scheres SHW. 2015. How cryo-EM is revolutionizing structural biology. Trends Biochem Sci 40:49–57. doi:10.1016/j.tibs.2014.10.005

Bartesaghi A, Lecumberry F, Sapiro G, Subramaniam S. 2012. Protein secondary structure determination by constrained single-particle cryo-electron tomography. Structure 20:2003–13. doi:10.1016/j.str.2012.10.016

Basanta B, Chowdhury S, Lander GC, Grotjahn DA. 2020. A guided approach for subtomogram averaging of challenging macromolecular assemblies. J Struct Biol X 4:100041. doi:10.1016/j.yjsbx.2020.100041

Bharat TAM, Russo CJ, Löwe J, Passmore LA, Scheres SHW. 2015. Advances in Single-Particle Electron Cryomicroscopy Structure Determination applied to Sub-tomogram Averaging. Structure 23:1743–1753. doi:10.1016/j.str.2015.06.026

Burnley T, Palmer CM, Winn M. 2017. Recent developments in the CCP-EM software suite. Acta Crystallogr Sect D, Struct Biol 73:469–477. doi:10.1107/S2059798317007859

Burt A, Cassidy CK, Ames P, Bacia-Verloop M, Baulard M, Huard K, Luthey-Schulten Z, Desfosses A, Stansfeld PJ, Margolin W, Parkinson JS, Gutsche I. 2020. Complete structure of the chemosensory array core signalling unit in an E. coli minicell strain. Nat Commun 11:743. doi:10.1038/s41467-020-14350-9

Castaño-Díez D. 2019. Dynamo: Walkthrough on GUI based tilt series alignment. https://wiki.dynamo.biozentrum.unibas.ch/w/index.php/Walkthrough_on_GUI_based_tilt_series_alignment

Castaño-Díez D, Kudryashev M, Arheit M, Stahlberg H. 2012. Dynamo: a flexible, user-friendly development tool for subtomogram averaging of cryo-EM data in high-performance computing environments. J Struct Biol 178:139–51. doi:10.1016/j.jsb.2011.12.017

Castaño-Díez D, Kudryashev M, Stahlberg H. 2017. Dynamo Catalogue: Geometrical tools and data management for particle picking in subtomogram averaging of cryo-electron tomograms. J Struct Biol 197:135–144. doi:10.1016/j.jsb.2016.06.005

Chen M, Bell JM, Shi X, Sun SY, Wang Z, Ludtke SJ. 2019. A complete data processing workflow for cryo-ET and subtomogram averaging. Nat Methods 16:1161–1168. doi:10.1038/s41592-019-0591-8

Chen M, Dai W, Sun SY, Jonasch D, He CY, Schmid MF, Chiu W, Ludtke SJ. 2017. Convolutional neural networks for automated annotation of cellular cryo-electron tomograms. Nat Methods 14:983–985. doi:10.1038/nmeth.4405

Ertel KJ, Benefield D, Castaño-Diez D, Pennington JG, Horswill M, den Boon JA, Otegui MS, Ahlquist P. 2017. Cryo-electron tomography reveals novel features of a viral RNA replication compartment. Elife 6. doi:10.7554/eLife.25940

Harris CR, Millman KJ, van der Walt SJ, Gommers R, Virtanen P, Cournapeau D, Wieser E, Taylor J, Berg S, Smith NJ, Kern R, Picus M, Hoyer S, van Kerkwijk MH, Brett M, Haldane A, Del Río JF, Wiebe M, Peterson P, Gérard-Marchant P, Sheppard K, Reddy T, Weckesser W, Abbasi H, Gohlke C, Oliphant TE. 2020. Array programming with NumPy. Nature 585:357–362. doi:10.1038/s41586-020-2649-2

Himes BA, Zhang P. 2018. emClarity: software for high-resolution cryo-electron tomography and subtomogram averaging. Nat Methods 15:955–961. doi:10.1038/s41592-018-0167-z

Kremer JR, Mastronarde DN, McIntosh JR. 1996. Computer visualization of three-dimensional image data using IMOD. J Struct Biol 116:71–6. doi:10.1006/jsbi.1996.0013

Leigh KE, Navarro PP, Scaramuzza S, Chen W, Zhang Y, Castaño-Díez D, Kudryashev M. 2019. Subtomogram averaging from cryo-electron tomograms. Methods Cell Biol 152:217–259. doi:10.1016/bs.mcb.2019.04.003

Moebel E, Martinez-Sanchez A, Larivière D, Fourmentin E, Ortitz J, Baumeister W, Kervrann C. 2021. Deep Learning Improves Macromolecules Localization and Identification in 3D Cellular Cryo-Electron Tomograms. bioRxiv. doi:10.1101/2020.04.15.042747

Nakane T, Kotecha A, Sente A, McMullan G, Masiulis S, Brown PMGE, Grigoras IT, Malinauskaite L, Malinauskas T, Miehling J, Uchański T, Yu L, Karia D, Pechnikova E V, de Jong E, Keizer J, Bischoff M, McCormack J, Tiemeijer P, Hardwick SW, Chirgadze DY, Murshudov G, Aricescu AR, Scheres SHW. 2020. Single-particle cryo-EM at atomic resolution. Nature 587:152–156. doi:10.1038/s41586-020-2829-0

Paszke A, Gross S, Massa F, Lerer A, Bradbury J, Chanan G, Killeen T, Lin Z, Gimelshein N, Antiga L, Desmaison A, Köpf A, Yang E, DeVito Z, Raison M, Tejani A, Chilamkurthy S, Steiner B, Fang L, Bai J, Chintala S. 2019. PyTorch: An Imperative Style, High-Performance Deep Learning Library.

Sanchez RM, Zhang Y, Chen W, Dietrich L, Kudryashev M. 2020. Subnanometer-resolution structure determination in situ by hybrid subtomogram averaging - single particle cryo-EM. Nat Commun 11:3709. doi:10.1038/s41467-020-17466-0

Schaffer M, Pfeffer S, Mahamid J, Kleindiek S, Laugks T, Albert S, Engel BD, Rummel A, Smith AJ, Baumeister W, Plitzko JM. 2019. A cryo-FIB lift-out technique enables molecular-resolution cryo-ET within native Caenorhabditis elegans tissue. Nat Methods 16:757–762. doi:10.1038/s41592-019-0497-5

Scheres SHW. 2012. RELION: implementation of a Bayesian approach to cryo-EM structure determination. J Struct Biol 180:519–30. doi:10.1016/j.jsb.2012.09.006

Schur FK. 2019. Toward high-resolution in situ structural biology with cryo-electron tomography and subtomogram averaging. Curr Opin Struct Biol 58:1–9. doi:10.1016/j.sbi.2019.03.018

Sofroniew N, Lambert T, Evans K, Winston P, Nunez-Iglesias J, Bokota G et al.. 2021. napari/napari: 0.4.3rc1. doi:10.5281/zenodo.4437554

Tegunov D, Cramer P. 2019. Real-time cryo-electron microscopy data preprocessing with Warp. Nat Methods 16:1146–1152. doi:10.1038/s41592-019-0580-y

Tegunov D, Xue L, Dienemann C, Cramer P, Mahamid J. 2021. Multi-particle cryo-EM refinement with M visualizes ribosome-antibiotic complex at 3.7 Å inside cells. bioRxiv.

ThermoFisher Scientific. 2020. Tomography. Software version 5.X. Release Notes. https://assets.thermofisher.com/TFS-Assets/MSD/Product-Updates/105074-TOMO-Release-notes.pdf

Turoňová B, Schur FKM, Wan W, Briggs JAG. 2017. Efficient 3D-CTF correction for cryo-electron tomography using NovaCTF improves subtomogram averaging resolution to 3.4Å. J Struct Biol 199:187–195. doi:10.1016/j.jsb.2017.07.007

Turoňová B, Sikora M, Schürmann C, Hagen WJH, Welsch S, Blanc FEC, von Bülow S, Gecht M, Bagola K, Hörner C, van Zandbergen G, Landry J, de Azevedo NTD, Mosalaganti S, Schwarz A, Covino R, Mühlebach MD, Hummer G, Krijnse Locker J, Beck M. 2020. In situ structural analysis of SARS-CoV-2 spike reveals flexibility mediated by three hinges. Science 370:203–208. doi:10.1126/science.abd5223

Urzhumtseva L, Urzhumtsev A. 2019. py_convrot: rotation conventions, to understand and to apply. J Appl Crystallogr 52:869–881. doi:10.1107/S1600576719007313

Virtanen P, Gommers R, Oliphant TE, Haberland M, Reddy T, Cournapeau D, Burovski E, Peterson P, Weckesser W, Bright J, van der Walt SJ, Brett M, Wilson J, Millman KJ, Mayorov N, Nelson ARJ, Jones E, Kern R, Larson E, Carey CJ, Polat İ, Feng Y, Moore EW, VanderPlas J, Laxalde D, Perktold J, Cimrman R, Henriksen I, Quintero EA, Harris CR, Archibald AM, Ribeiro AH, Pedregosa F, van Mulbregt P, SciPy 1.0 Contributors. 2020. SciPy 1.0: fundamental algorithms for scientific computing in Python. Nat Methods 17:261–272. doi:10.1038/s41592-019-0686-2

Wan W, Briggs JAG. 2016. Cryo-Electron Tomography and Subtomogram Averaging. Methods Enzymol 579:329–67. doi:10.1016/bs.mie.2016.04.014

