## Supplementary Figures for "Tools enabling flexible approaches to high-resolution subtomogram averaging"

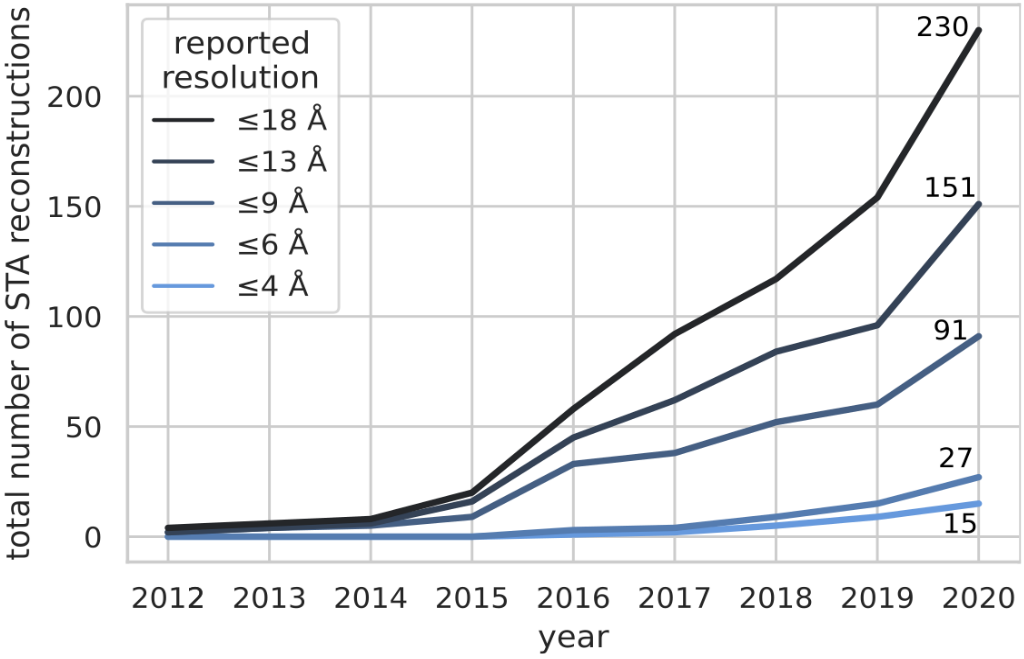


**Figure 1, figure supplement 1**: The number of subtomogram averaging reconstructions with different reported resolution values in the EMDB

```python
import numpy as np
import mrcfile
import starfile
from eulerangles import euler2matrix, invert_rotation_matrices
import napari

## Get position, orientation and imaging data to be visualised
# Set file paths
particle_file = 'particles_10.00Apx.star'
volume_file = 'TS_01.mrc_10.00Apx.mrc'

# Read data into memory
star = starfile.read(particle_file)
volume = mrcfile.open(volume_file).data

# Isolate the subset of particles which correspond to the tomogram
subset = star.loc[star['rlnMicrographName'] == volume]

# Get Euler angles and positions from dataframe as NumPy arrays
eulers = subset[[f'rlnAngle{angle}' for angle in ('Rot', 'Tilt', 'Psi')]].to_numpy()
coords = subset[[f'rlnCoordinate{axis}' for axis in 'XYZ']].to_numpy()

# Derive rotation matrices from euler angles & invert to yield the active transformation
rotation_matrices = euler2matrix(eulers, axes='zyz', intrinsic=True,
right_handed_rotation=True)
rotation_matrices = invert_rotation_matrices(rotation_matrices)

# Calculate rotated unit z vectors (Rv = v')
unit_z = np.array([0, 0, 1]).reshape((3, 1))
rotated_unit_z = rotation_matrices @ unit_z

## Set up napari visualisation
# Imaging data axis order is zyx, our coords are xyz - reorder coords and vectors
coords_napari = coords[:, ::-1]
rotated_unit_z_napari = rotated_unit_z.squeeze()[:, ::-1]

# Set up vectors layer data - https://napari.org/tutorials/fundamentals/vectors.html
vec_shape = (coords.shape[0], 2, 3)
vectors_napari = np.empty(vec_shape)
vectors_napari[:, 0, :] = coords_napari
vectors_napari[:, 1, :] = rotated_unit_z_napari

# Create visualisation
with napari.gui_qt():
    viewer = napari.Viewer(ndisplay=3)
    vol_layer = viewer.add_image(volume)
    pos_layer = viewer.add_points(coords_napari, size=4.5, face_color='black')
    vec_layer = viewer.add_vectors(vectors_napari, length=10, edge_color='cornflowerblue')
```

**Figure 2, figure supplement 1**: The source code for generating the scene depicted in Figure 2, combining *starfile* and *eulerangles* with existing tools in the scientific Python ecosystem.
